# A method for rapid nanobody screening with no bias of the library diversity

**DOI:** 10.1101/2023.02.15.528753

**Authors:** Zhiqing Tao, Xiaoling Zhao, Huan Wang, Juan Zhang, Guosheng Jiang, Bin Yu, Yihao Chen, Mingjun Zhu, Junli Long, Lei Yin, Xu Zhang, Maili Liu, Lichun He

## Abstract

Nanobody refers to the variable domain of heavy-chain-only antibodies. The distinctive advantages of nanobodies including small size, feasible expression in *Escherichia coli* (*E. coli*), and superior stability make them promising tools for applications in scientific research and therapies. So far, the screening and expression of nanobodies are mainly following similar methods used for conventional antibodies, suffering from amplification-caused losses of the diversity of libraries and requirements of subcloning of interests into the expression vector. Here, based on the unique properties of nanobodies, we developed an integrated method to screen and express nanobodies simultaneously with no bias of the library diversity. The library of nanobodies was cloned and secretively expressed into the culture medium. Target specifical binding nanobodies were isolated through 1-3 rounds of dilution and regrown steps in a way following the Poisson distribution to ensure no positive clones were dismissed, while the population of positive clones increased by more than 10 folds upon each round of dilution. Ultimately, 5 nanobodies against the death domain receptor 5 (DR5) and 5 nanobodies against the *Pyrococcus furiosus* (*Pfu*) DNA polymerase were produced directly out of their immunized libraries, respectively. Additionally, our approach allowed nanobody screening even without any specialized instruments/devices, demonstrating general applicability in the routine production of monoclonal nanobodies for diverse biomedical applications.

## Introduction

Single domain antibody also named nanobody, refers to the variable domain from the heavy chain only antibody (VHH) ^*1*^. Nanobody emerges as an alternative to the traditional antibody as it has several unique properties, such as high solubility, excellent stability, and small size (10-15 kDa) ^*2*^. Unlike the normal monoclonal antibody, nanobody can be expressed in *E. coli* with a high yield. Fusing the N terminus of a nanobody with a secretion signal peptide can lead to the secretion of the nanobody into the periplasmic space and the extracellular milieu through the type II secretion system ^3^ or some other unspecific release ways ^4^. Moreover, nanobodies could specifically bind the antigen with an affinity in the nanomolar range ^5^. The lack of light chains facilitates nanobodies for the construction of bispecific antibodies as well as conjugations with E3 ubiquitin ligases for specific degradation of target proteins ^6^. Owing to these properties of the nanobody, diverse biomedical and scientific applications of nanobody also include the usage of nanobody in hot start polymerase chain reactions (PCRs) and chimeric antigen receptor T cell (CAR-T) therapies ^7,8^. Nevertheless, the isolation of nanobodies mainly follows methods used for screening traditional antibodies such as phage display, cell surface display, and ribosome display ^9^. These display methods are usually labour-intensive, involving separated screening and producing vector systems. The isolation of target-specific monoclonal antibodies usually takes months. Furthermore, the commonly used phage display method has been reported to suffer from the loss of library diversity during the amplification procedure ^10,11^. The amplification of the screening libraries, which is a critical step for phage display, diminishes the diversity of libraries and reduces the number of different clones binding with the antigen specifically, hindering the identification of antibodies with different binding epitopes and affinities. One of the reasons is the competition of the binding between antibodies and the target molecule, which reduces the monoclonal antibodies with lower affinities. Other factors rather than the binding affinity also contribute to the loss of diversities of the library. Different peptides are presented unequally by phages ^12^. Moreover, a single phage particle infects bacteria and then secrets more than 1,000 copies of phage, which greatly enriches phages with growth advantage during any of the amplification procedures ^10^. By contrast, several recent studies highlighted the importance of the diversity of antibodies. Low- or moderate-affinity rather than high-affinity antibodies deliver greater activity, even when they share overlapping binding epitopes with the high-affinity antibody^13-15^. Thus, the development of new methods for antibody discoveries is needed to overcome the problem of the convergence of the library.

Here, we presented a rapid and no-biased approach to generate monoclonal nanobodies against selected targets. Two immunized libraries against the Death receptor 5 (DR5) and the Pyrococcus furiosus (*Pfu*) DNA polymerase were secretively expressed into the culture medium respectively. With the dilution and regrowth of *E. coli* transformed with each immunized library, populations of positive nanobody clones were enriched more than 10-fold per round with no loss of the library diversity. The target specifical monoclonal nanobodies were then isolated and purified directly from the culture medium. Moreover, this integrated way of screening and expression of nanobodies works even without any specialized instruments, providing a robust, low-cost and no-biased approach to generating monoclonal nanobodies against selected targets.

## Results

### Secretion of nanobodies into the culture medium

*E. coli* has been used as one of the most suitable hosts for the production of recombinant proteins, such as nanobodies and antibody fragments. However, nanobodies expressed in the cytosol of *E. coli* face several problems including the formation of insoluble inclusion bodies, and the inability to form disulfide bonds correctly ^16^. Both will ultimately lead to toxicity to the host cell ^17^. As data from our lab showed several proteins could be secreted into the culture medium by fusing it with the pelB signal peptide (Figure S1 HdeA, Spy, Im7 secretion). We are wondering if nanobodies could be secreted into the culture medium to avoid the aforementioned toxic problems. For this purpose, we constructed expression vectors by fusing the pelB signal peptide and 6× His-tags on the N-terminus of nanobodies from a commercial naive library (Qinhe Life Science Ltd.) (Figure 1B). As we expected, the preliminary experiments revealed all 5 randomly chosen nanobodies were secreted into the culture medium with a clear band on the SDS PAGE gel (Figure 1C), proving the feasibility of secretion expression of nanobodies by *E. coli*. The secretion of nanobody into the culture medium not only enables us to screen the monoclonal nanobody in a similar way as the hybridoma technology but also allows us to integrate the screening and purification of nanobodies by using the same expression vector and host cells, which will greatly reduce the time needed for the discovery of specific binding nanobodies.

**Figure. 1:**
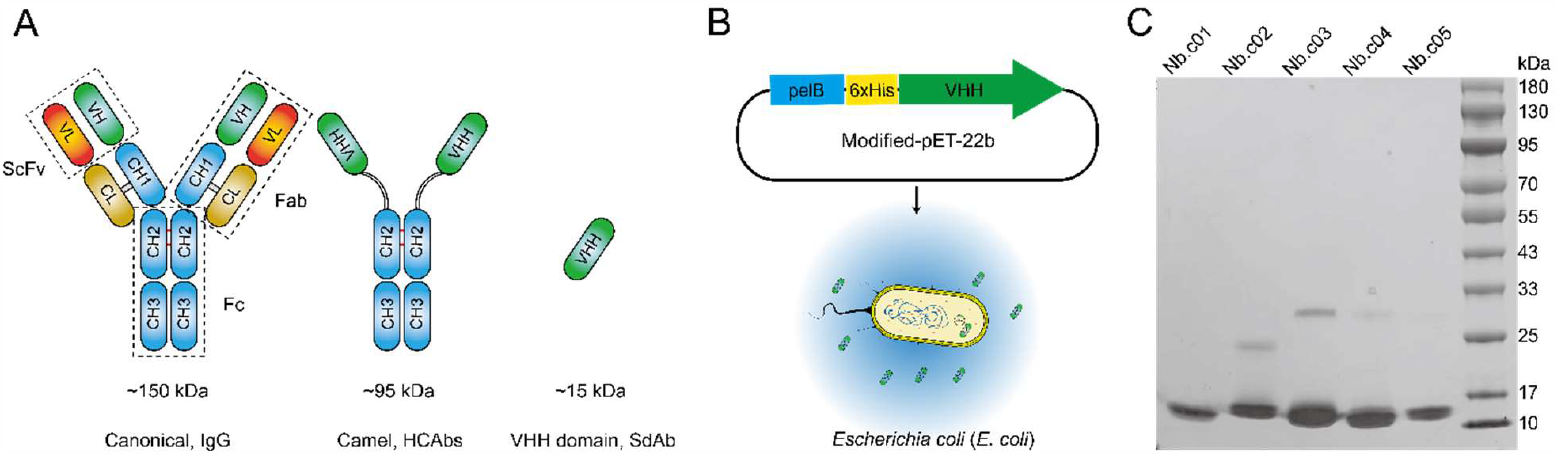
Unique features of nanobodies compared to conventional antibodies (A) Schematic structures of the conventional antibody, the heavy chain only antibody from camel serum, and the variable fragment of heavy chain antibodies (VHH) also named as nanobody or single domain antibody (SdAb). (B). Construction and modification of the pET22b vector for expression and secretion of nanobodies by *E. coli*. (C) SDS-PAGE analysis of various nanobodies secreted into the *E. coli* culture medium. 10 times concentrated supernatant of the cell culture medium was applied for SDS-PAGE analysis.

### New strategy for direct isolation of specific nanobodies with simple dilution and regrown cycles

The numbers of antigen-specific immunoglobulin G (IgG)-secreting B cells in the peripheral blood range from the order of 10^1-4^ per 10^6^ cells after a single immunization ^18-20^. Thus, directly following the hybridoma technology by diluting an immunized nanobody library expressing *E. coli* into single cell per well to perform screening for specific monoclonal nanobodies faces the problem of low throughput ^21^. To solve this problem, we designed a new strategy to mix 10^n^ clones per well. In this way, a single 96-well plate could reach a screening throughput of 10^n+2^. For the convenience of calculation, 96 wells were approximated as 100 wells here and in the following text. Once the well with positive clones was identified, a division of m clones was taken out, diluted, and re-distributed equally into a new 96-well plate for the next round of isolating the specific monoclonal nanobody. Since the mixture of *E. coli* clones in each well could be considered as complete homogenous materials, the number of positive clones in the division (m) taken out from the positive well follows the Poisson distribution (equation 1),

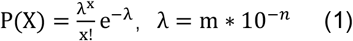

where P(X) was the probability of taking X positive clones; λ specified the expected value of the number of positive clones in the division (m), which equaled the product of the number of clones in the division (m) and the probability of taking one positive clone (10^-n^); e ≈ 2.718 was the base of the natural logarithm. The probability of at least one positive clone was presented in the division (m) for diluting, distributing, and regrowing for the next round of screening could be calculated with equation (2).

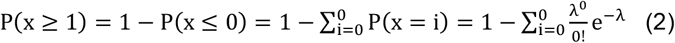

The population of positive clones will be enriched by a factor of ^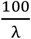^ over each round of isolation. Accordingly, the probability of at least one positive clone was taken in the division (m) from the identified positive well was 0.993262, 0.999955, and 0.999999, when λ equaled 5, 10, and 20 respectively. The population of positive clones will be enriched by a factor of 20, 10, and 5 respectively in the next round of isolation (Figure S2). A higher value of λ helped to avoid the dismission of positive clones in the process of screening. However, considering the enriching factor of positive clones over each round of screening, a λ value of 10 was used in the following study.

### Assessing the performance of the new strategy with two immunized libraries

To assess the performance of the new strategy above, we first immunized two alpacas with purified antigen proteins DR5 and *Pfu* DNA polymerase respectively. IgG titers assayed by ELISA confirmed the generation of specific camel antibodies against both antigen proteins in the serum of immunized animals (Figure S3). Total mRNA was extracted from lymphocytes according to the protocol of the LeukoLOCK™ Total RNA Isolation Kit. The VHH gene segments from two immunized cDNA libraries were amplified by nested PCR and then cloned into our modified pET-22b secretion vector. For *Pfu* DNA polymerase nanobodies, upon the transformation of *E. coli* with constructed nanobody encoding vectors, the first round of screening was performed with 10^3^ clones inoculated in 2 mL TB culture medium in each well of a 96 deep-well plate (A_1_ plate). All wells in the A_1_plate were induced with 0.5mM IPTG for 12 hours and centrifuged to collect supernatants for ELISA assay to identify the well(s) with positive clones. Since all secreted nanobodies had His-tags constructed on the N-terminus, the ELISA plate was then incubated with an anti-His-tag mouse monoclonal antibody and a secondary anti-mouse antibody conjugated horse radish peroxidase (HRP) to detect specific bound nanobodies. The screening results were shown in Figure 3A-D. Several wells showed OD_450_ values large than 1.0, indicating *E. coli* clones expressing specific nanobodies against the *Pfu* DNA polymerase were presented in these well. The probability of certain positive clones (p_i_) was ∼1‰. Cells in the positive well of the A_1_ plate were then resuspended to determine the cell number by using the standard curve of the cell number and the value of OD_600_ (Figure S4). A division of ∼10^4^ clones was taken out from the positive well to dilute and aliquot equally into a new 96 deep-well plate (A_2_) for regrowing. The expected value (λ) of the number of a certain positive clone presenting in the division was 10 according to equation 1. The probability of positive clones taken into the division from the identified positive well was 99.995% according to equation 2. Therefore, hardly any positive clone would be dismissed with our new strategy. As these ∼10^4^ clones taken from the positive well were aliquoted evenly into the A_2_ plate, each well will have ∼100 clones. The probability of a certain positive clone presenting in one well increased from 1‰ to 1% (Figure 2). With 2 rounds of screening, the number of clones in each well was ∼ 10 (Figure 3C). A simple diluting and spreading of cells from the positive well on the LB agar plate with ampicillin were performed to get single cell clones of *E. coli*. The single cell clones were then inoculated into another 96 deep-well plate (A_3_) for the final round of screening to get the specific monoclonal nanobodies (Figure 2, Figure 3D). In the end, 6 nanobodies against *Pfu* DNA polymerase were isolated, of which 5 different nanobodies were identified according to the calculated genetic distance ^22^ (Figure 3H). For DR5 nanobodies, the first round of screening with 10^2^ clones inoculated per well showed all positive signals, indicating the population of DR5-specific immunoglobulin G (IgG)-secreting B cells was large. We then tried to isolate the DR5-specific monoclonal nanobody by picking single cell clones from the LB plate to inoculate in the 96 deep-well plate with one clone per well. 22 out of 96 clones revealed positive signals from the ELISA assay, showing the high immunogenicity of DR5 (Figure 3E). 10 clones from the positive wells were picked and sent for sequencing. 5 different nanobodies were identified according to the calculated genetic distance ^22^ (Figure 3G). Nevertheless, we would like to point out more different nanobodies against both *Pfu* DNA polymerase and DR5 could be isolated if needed. Both isolated *Pfu* DNA polymerase and DR5 specific nanobody expressing *E. coli* strains were used directly for productions of monoclonal nanobodies. After one step of purification by the Ni-NTA column, the purity of nanobodies reached >95% by SDS-PAGE (Figure S5). This integrated screening and purification of target-specific monoclonal nanobodies did not need to graft the gene of interest into a new expression vector, simplifying the discovery of nanobodies.

**Figure. 2:**
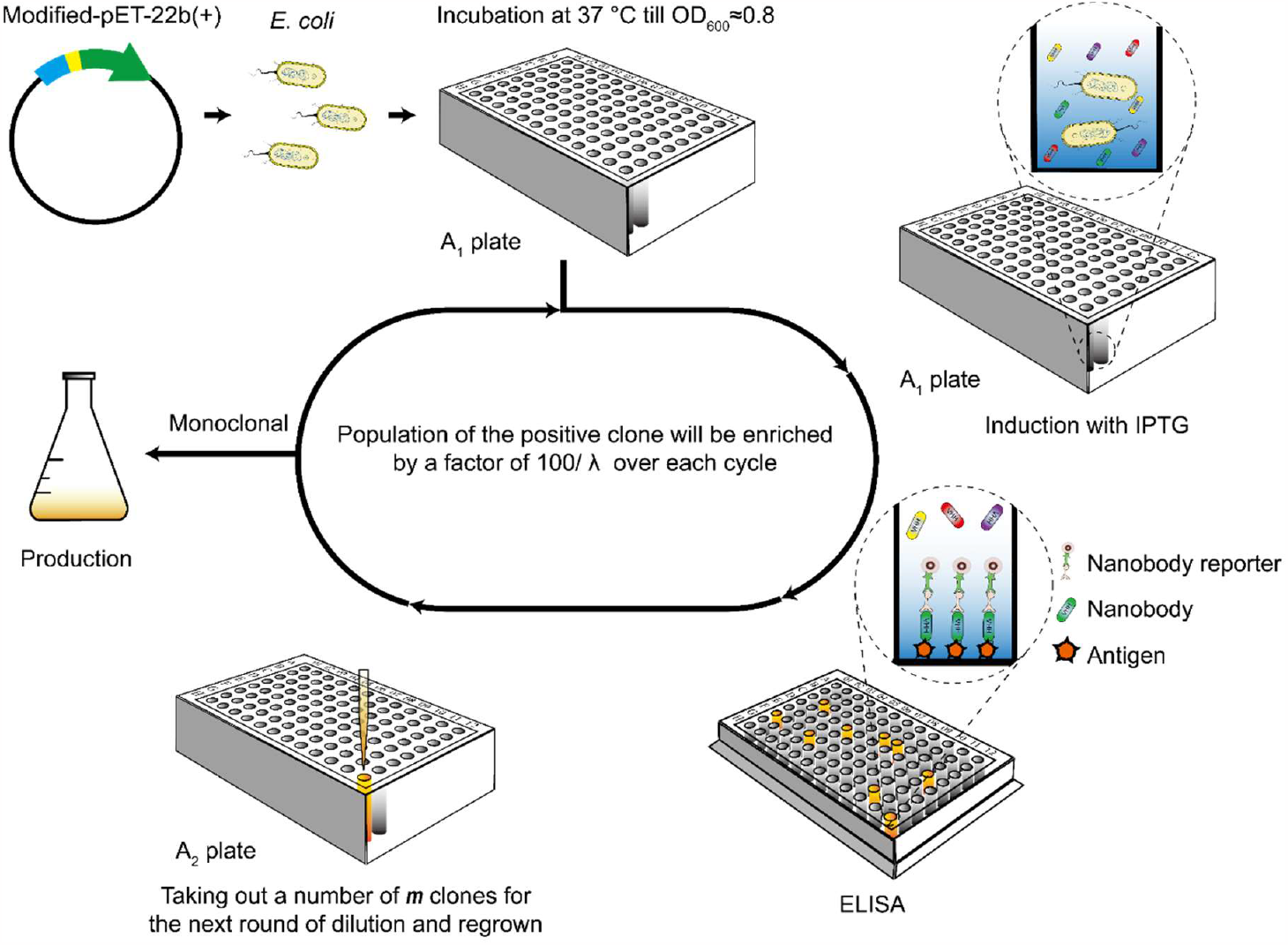
Scheme of integrated isolation and expression of the specific binding monoclonal nanobody *via* repeated dilution, distribution, and regrown cycles following the rule of Poisson distribution to avoid the dismission of the positive clones while enriching the population of positive clones.

**Figure. 3:**
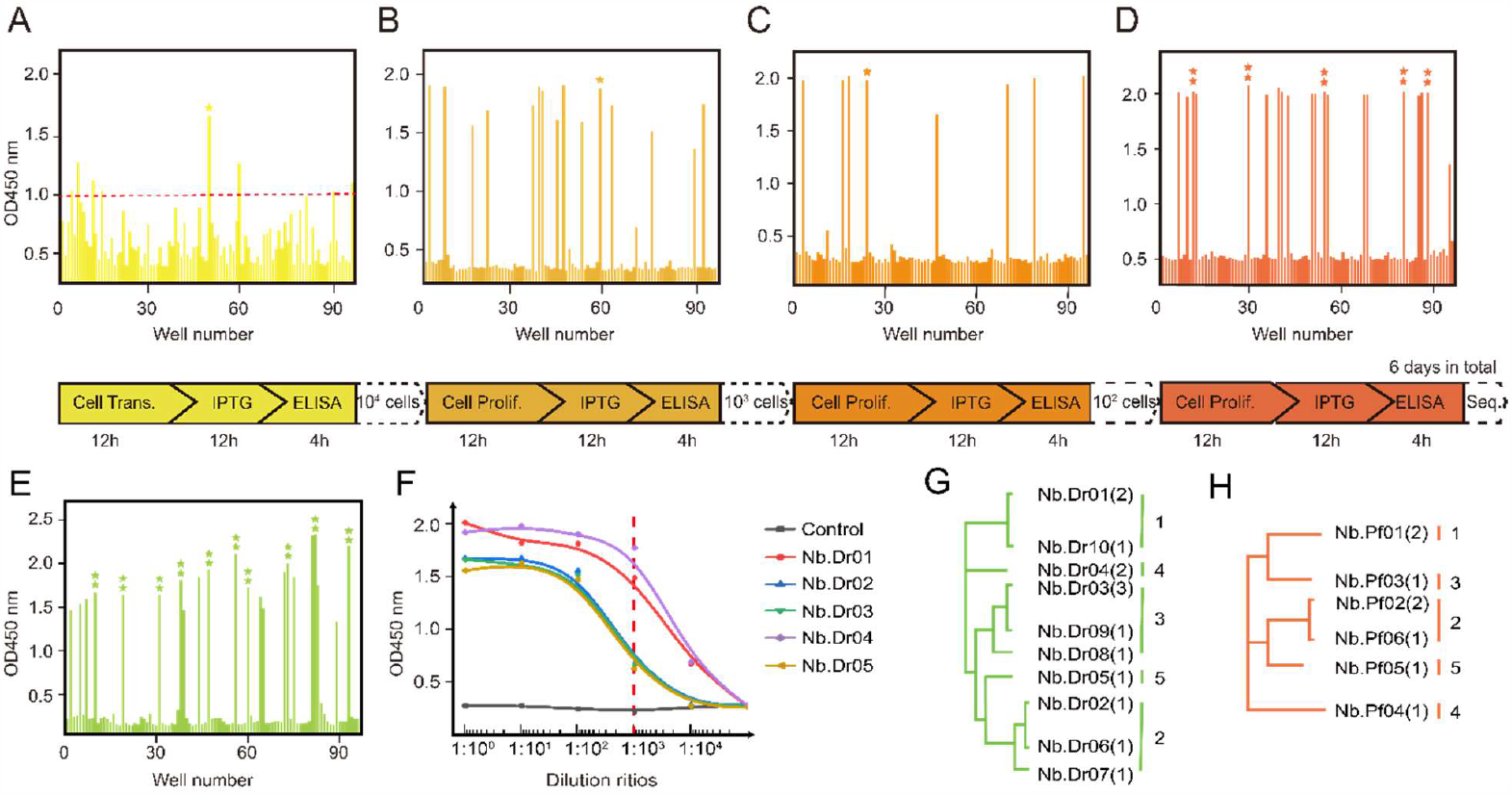
Isolation of nanobodies against *Pfu* DNA polymerase and DR5. ELISA results of supernatants of nanobody culture media. (A-C). Supernatants of ∼1000 (A), ∼100 (B), ∼10 (C) different nanobody expressing clones were monitored against *Pfu* DNA polymerase by ELISA. * indicated the positive well picked for the next round of screening. (D-E) Supernatants of single nanobody expressing clones were detected against *Pfu* DNA polymerase (D) and DR5 (E) by ELISA, respectively. ⁑ indicated the single clone isolated for sequencing. f. Supernatants of different DR5 nanobody expressing clones were monitored by ELISA upon dilution with the supernatant of *E. coli* transformed with a naive library of nanobodies by ratios of 1:10^1^, 1:10^2^, 1:10^3^, and 1:10^4^ respectively. (G-H) Phylogenetic tree showing genetic distances between different nanobodies of *Pfu* DNA polymerase (H) or DR5 (G).

### High throughput of the new strategy for isolating nanobodies

The screening throughput of our strategy was dependent on the number of clones inoculated per well. The signal of positive clones was highly related to its secretion level in the culture medium. For the nanobodies against *Pfu* DNA polymerase, we inoculated 10^3^ clones per well, showing a screening throughput of 10^5^ for a single plate. However, nanobodies against DR5 were isolated with the inoculation of a single clone per well. To demonstrate the screening throughput for DR5 nanobodies, we diluted the supernatant of each of five isolated DR5 specific nanobodies with the supernatant from *E. coli* transformed with a naive library of nanobodies by ratios of 1:10^1^, 1:10^2^, 1:10^3^, and 1:10^4^ respectively. All five nanobodies showed clear positive signals at the dilution ratio of 1:10^3^. Two DR5-specific nanobodies even showed a positive signal upon dilution by the ratio of 1:10^4^, demonstrating secretion levels of naturally occurred nanobodies in our system were fairly high to support a screening throughput of 10^5^-10^6^ for a single 96-well plate (Figure. 3F).

### A broad range of binding affinities of isolated nanobodies against target proteins

We performed isothermal titration calorimetry (ITC) assays to access the binding affinities of isolated nanobodies. DR5, *Pfu* DNA polymerase, and their corresponding isolated nanobodies were purified and dialyzed in the same buffer 20 mM HEPES, 150 mM NaCl, pH 8.0 at 4 °C overnight. The results showed binding affinities of five isolated DR5 nanobodies with DR5 ranged from 18 μM to 54 pM and the binding affinities of four isolated *Pfu* nanobodies with *Pfu* DNA polymerase ranged from 33 μM to 35 pM (Table 1, Figure 4 and Figure S6), confirming our developed strategy had with no bias of high affinity antibodies and could be applied for isolating nanobodies with a broad range of binding affinities with target proteins.

**Table 1:** Affinity data for DR5 and *Pfu* with their respective nanobodies.

| Antigen | Nanobody | K<sub>D</sub> [M] |
| --- | --- | --- |
| DR5 | Nb.Dr01 | $(5.4 \pm 0.4) \times 10^{-11}$ |
| | Nb.Dr02 | $(9.5 \pm 6.3) \times 10^{-10}$ |
| | Nb.Dr03 | $(1.0 \pm 0.1) \times 10^{-7}$ |
| | Nb.Dr04 | $(12.0 \pm 1.0) \times 10^{-7}$ |
| | Nb.Dr05 | $(1.8 \pm 0.3) \times 10^{-5}$ |
| *Pfu* DNA<br>polymerase | Nb.Pf01 | $(3.5 \pm 0.1) \times 10^{-11}$ |
| | Nb.Pf02 | $(3.5 \pm 1.0) \times 10^{-8}$ |
| | Nb.Pf03 | $(3.3 \pm 2.1) \times 10^{-5}$ |
| | Nb.Pf04 | $(5.9 \pm 2.4) \times 10^{-8}$ |
|  | Nb.Pf05 | Data not available |

**Figure 4:**
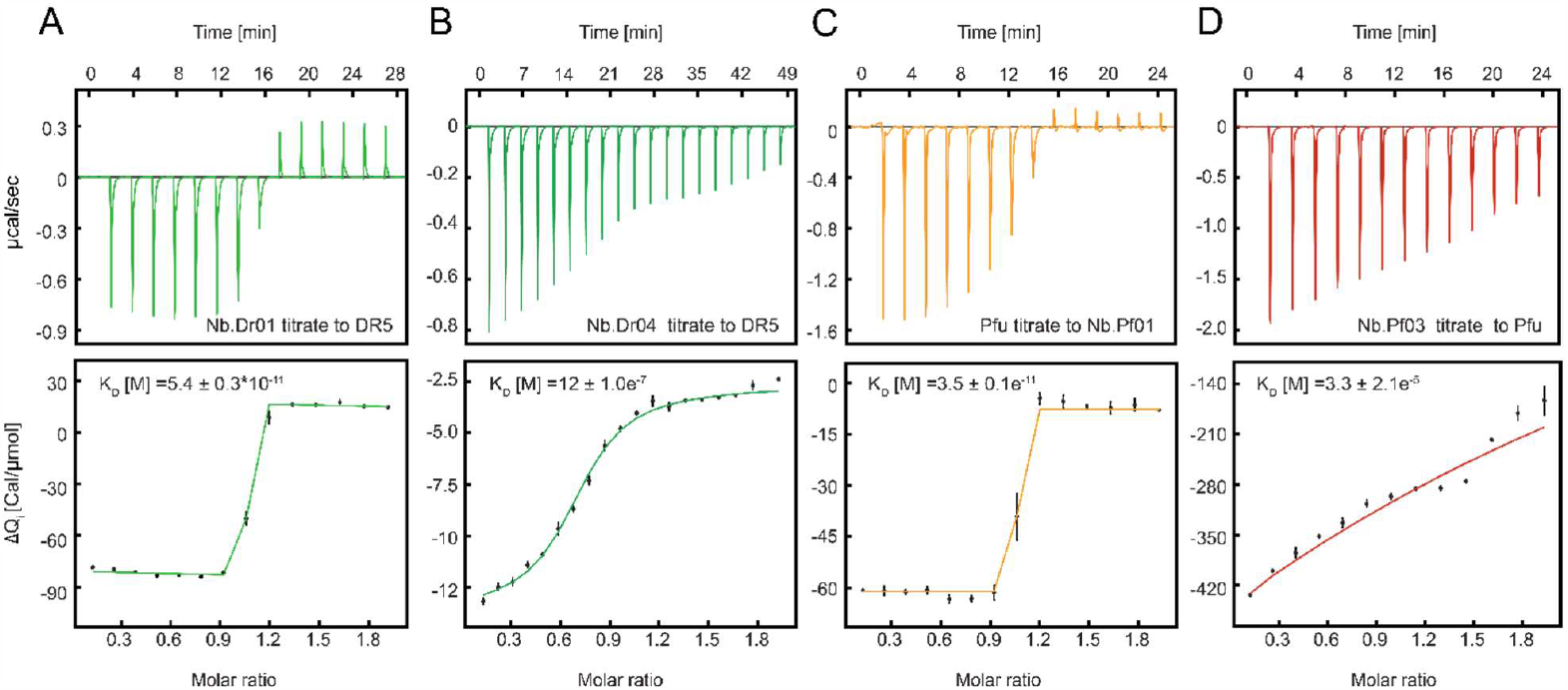
Determination of the binding affinities of isolated nanobodies against their target proteins. Representative results from ITC measurements of two monoclonal nanobodies against DR5 protein (A-B) and two monoclonal nanobodies against *Pfu* DNA polymerase (C-D). ITC measurements of all isolated nanobodies against their target proteins are shown in Figure S6.

### Binding epitopes of isolated nanobodies on *Pfu* DNA polymerase

To demonstrate the diversities of isolated nanobodies, we further probed if the binding epitopes of different nanobodies on *Pfu* DNA polymerase overlapped with each other *via* NMR spectroscopy. For each of the isolated *Pfu* nanobodies, both ^15^N labeled and unlabeled samples were expressed and purified. The ^15^N-^1^H HSQC spectrum of each isolated nanobody was first measured in the presence and absence of *Pfu* DNA polymerase. Upon adding *Pfu* DNA polymerase, the formation of the nanobody and *Pfu* DNA polymerase complex caused a shorter T2 relaxation time and the consequent peak intensity reduction of the labeled nanobody in the spectrum. (Figure 5A and Figure S7). Then ^15^N-^1^H HSQC spectra of each *Pfu* nanobody were recorded in the presence of *Pfu* DNA polymerase and an another different unlabeled nanobody. If two nanobodies shared overlapping binding sites on *Pfu* DNA polymerase, the addition of a second competitive unlabeled nanobody would release the ^15^N labeled nanobody from *Pfu* DNA polymerase and restore the peaking intensity of the ^15^N labeled nanobody. If two nanobodies had separated binding sites on *Pfu* DNA polymerase, the addition of a second non-competitive unlabeled nanobody would have little effect on the peaking intensity of the ^15^N labeled nanobody. In this way, the overall competitive status of all five nanobodies in binding with *Pfu* DNA polymerase was measured and summarized in Figure 5C. To connect the binding epitopes of the nanobody with the functional site of *Pfu* DNA polymerase, the inhibition of DNA polymerase activity of *Pfu* DNA polymerase by different nanobodies was accessed. (Figure 5D and Figure S8). Single-stranded DNA incubation assay was performed in the presence of each individual *Pfu* nanobody and the SYBR green I dye, which was used as an indicator of the final amount of double-strand DNA. The results showed the addition of either Nb.Pf02 or Nb.Pf04 strongly inhibited the polymerase activity of *Pfu* DNA polymerase, indicating that Nb.Pf02 and Nb.Pf04 bind to active sites of the polymerase domain of *Pfu* (Figure 5D). According to the mutual competitive relationship, the cartoon representation of binding sites of all five *Pfu* nanobodies was plotted in Figure 5B, with the binding site of Nb.Pf02 on the *Pfu* DNA polymerase as the reference.

**Figure 5:**
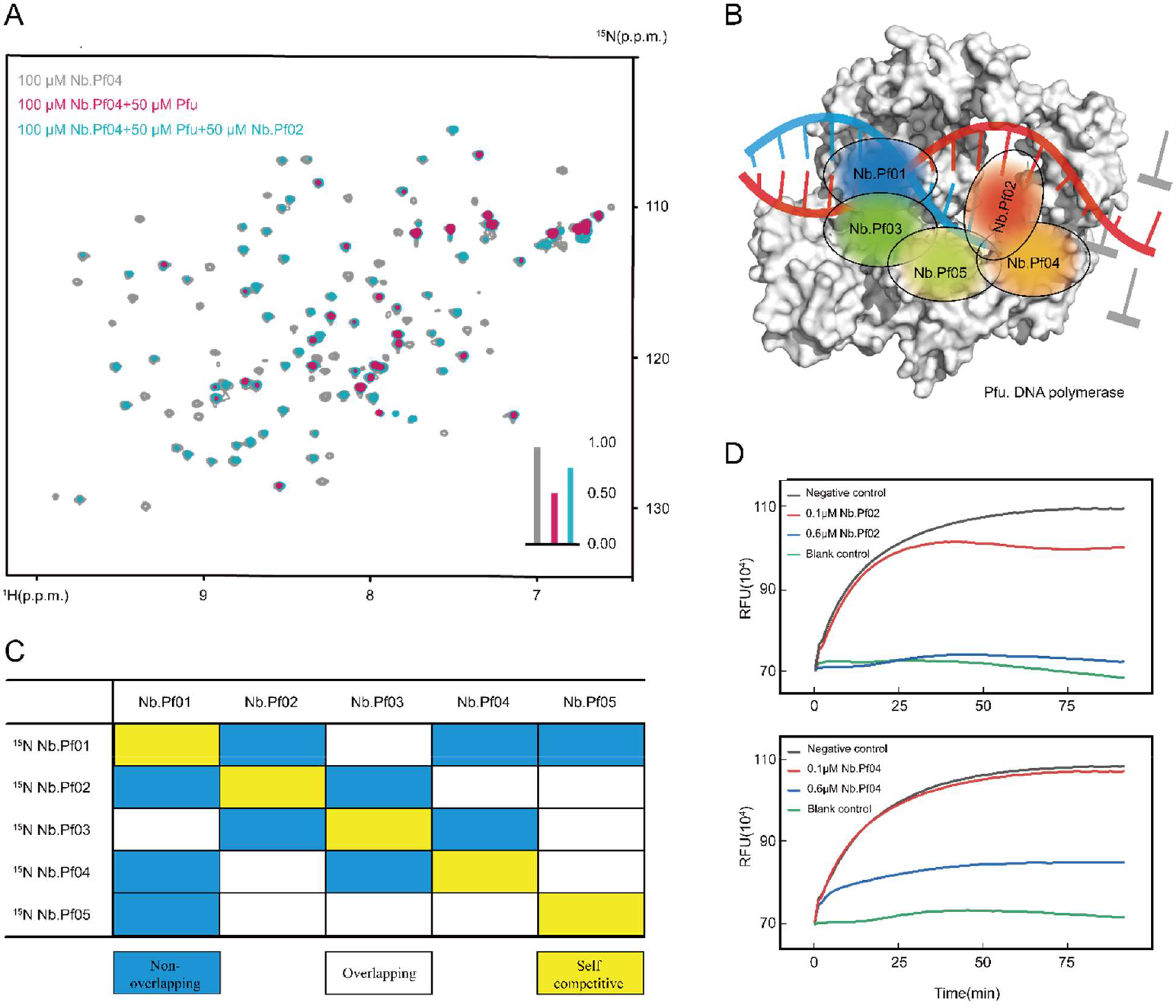
Characterization of the isolated nanobodies against *Pfu* DNA polymerase. A. ^15^N-^1^H HSQC spectra of 100 μM Nb. Pf04 in the presence (magenta) and absence (grey) 50 μM *Pfu* DNA polymerase. Addition of 50 μM unlabeled Nb.Pf02 partially restored the peak intensity of 100 μM Nb. Pf04 with 50 μM *Pfu* DNA polymerase (cyan). All spectra were recorded at 25 °C on a Bruker Avance-600 spectrometer. Data were processed in the same way and displayed with the same contour level. B. Cartoon representation of binding sites of all five nanobodies on *Pfu* DNA polymerase. C. A table summarized the overall competitive status of all five nanobodies upon interacting with *Pfu* DNA polymerase. D. The polymerase activity of *Pfu* DNA polymerase was blocked by either Nb. Pf04 or Nb.Pf02 monoclonal nanobody. The green line represented the blank group with no *Pfu* DNA polymerase. The black line represented the control group with *Pfu* DNA polymerase but in the absence of any nanobody. The red and blue lines represent experimental groups with *Pfu* DNA polymerase and a final concentration of 0.1 μM (red) and 0.6 μM (blue) of the isolated monoclonal nanobody, respectively.

## Discussion

Quick isolation of target-specific nanobodies is crucial for both research and therapies. Instead of the widely used screening methods, herein, we developed a method for the discovery of nanobodies from aspects of both theories and practices. Taking advantage of the small molecular weight and the feasibility of nanobodies to be expressed and secreted into the culture media by *E. coli*, our integrated way of screening and expression of specific nanobodies eliminated the step of subcloning of nanobodies into an expression vector. The direct work needed, including cDNA generation (0.5 d), nest PCR (0.5 d), transformation of vectors into *E. coli* (1 d), screening of nanobodies (4-5 d), and production of nanobodies (2 d). The entire work could be done in 8-9 days after the immune response was generated. Thus, this developed method facilitated reliable, rapid, and integrated screening and production of specific monoclonal nanobodies. Furthermore, this approach allowed nanobody screening at a low cost and worked even without any specialized instruments/devices. In case of limited access to the plate reader, our approach of isolating monoclonal antibodies could still be carried on by identifying positive clones with the naked eye. This is particularly helpful for laboratories in startup companies or labs in developing countries. We demonstrated the highest screening throughput of our method was ∼10^6^ per 96-well plate. Given the frequency of antigen-specific IgG secreting B cells often ranged from 10 to 1000 per 10^6^ cells after the immunization ^23^. Our approach is well suited for the discovery of monoclonal nanobody reagents from the immunized nanobody library. With our modified pET-22b vector, nanobodies are secreted and identified directly from the supernatant of *E. coli* culture media. The isolated target-specific nanobody could be directly used for fermentation to produce nanobodies on a large scale, greatly simplifying the process of production of new nanobodies from scratch.

In comparison with the commonly used phage display method, our approach avoids the problem of artificial loss of library diversity. Different phage clones have different growth rates in the common pool of bacteria ^10^. During the process of multiple rounds of panning and amplifying of phages, the gene of antibodies carried by the slow-growing phages will be lost ^11^. This decrease in library diversity may not be a problem for targets with a single binding site, but it could severely limit the number of useful antibodies identified for targets with different binding epitopes for antibodies. Our approach does not require panning and amplifying of the libraries; therefore, the diversity could be maintained, enabling the identification of novel ligands. Another advantage of our approach is its general applicability: the developed method enables the quick production of a comprehensive repertoire of *Pfu* DNA polymerase and DR5 binding nanobodies. For both DR5 and *Pfu* DNA polymerase, the isolated nanobodies show a wide range of binding affinity from tens of μM to pM, spanning up to 7 orders of magnitude, demonstrating the wide applicability of our methods for isolating nanobodies with a broad range of binding affinities.

In conclusion, our method was shown to enable rapid production of nanobodies with a few rounds of dilution and regrown of nanobody library transformed *E. coli* in 96-well plates. The advantages of the integrated screen and expression procedure, low-equipment dependency, and causing no loss of library diversity convince us our approach has general applicability for nanobody discoveries. Taking into account of advantages of nanobodies: nanobodies can be easily humanized for application in drug development and clinical therapies. They are much smaller than conventional antibodies and could be fused with each other to generate bivalent or multivalent nanobodies ^24^. The excellent chemical and thermal stability of nanobodies also promote their applications in antibody-drug conjugate (ADC) and proteolysis-targeting chimaeras (PROTAC) drugs ^25^. We envision that our method for nanobody discovery can be widely implemented in academia and industry to generate nanobody reagents suitable for various applications.

## Materials and Methods

### Construction of nanobody immune library

The immunization of llama was performed according to the published protocol ^26^. In short, a 3-year-old and a 4-year-old female llama were immunized by six weekly injections of 0.3 mg recombinant DR5 and *Pfu* DNA polymerase with equal volume of GERBU Adjuvant (GERBU Biotechnik GmbH), respectively. 10 mL of peripheral blood was collected as control prior to the immunization of each llama. 15 mL of peripheral blood was collected after every injection of the antigen protein mixed with GERBU Adjuvant. Serial dilutions of the pre-immune and immune serum were applied for ELISA to confirmation the generation of antigen specific nanobodies. The blood collected following the final immunization was used for extractions of total RNA with the commercial LeukoLOCK™ Total RNA Isolation Kit (Thermo Fisher). cDNA was obtained by the reverse transcription of RNA using the All-in-One 1st cDNA Synthesis MasterMix kit (Swiss Affinibody LifeScience AG). The VHH gene segments were amplified by nested PCR using CALL001, CALL002 ^27^, VHH-EcoR I-For and VHH-HindIII-Rev primers (Table 2) and cloned into a modified pET-22b vector.

**Table 2:**
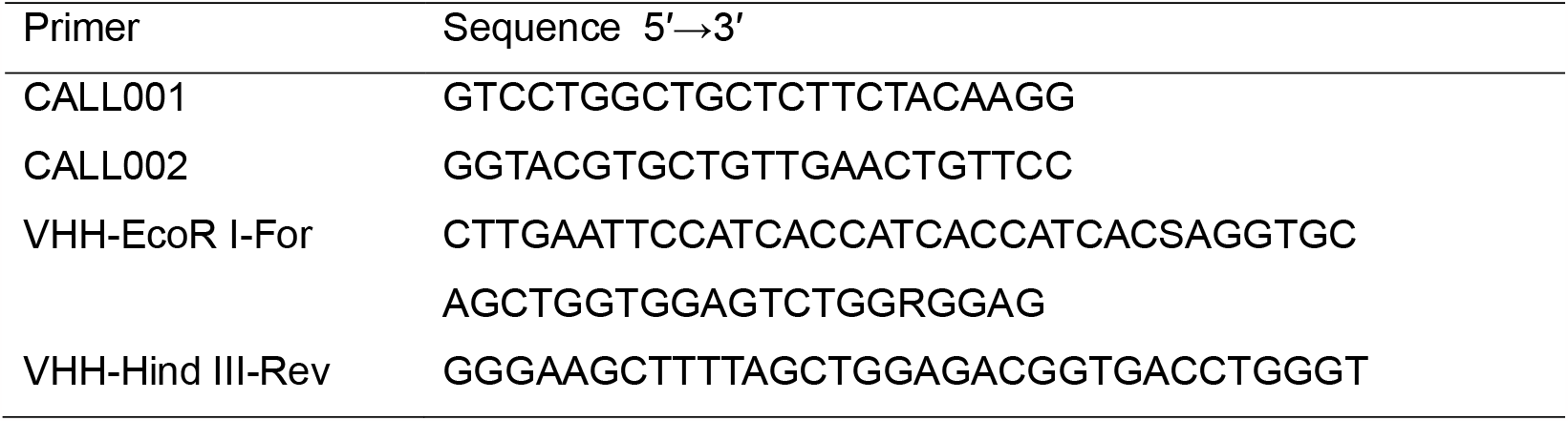
Primers for nested PCR of VHH genes.

### Enzyme-linked immunosorbent assays (ELISA)

96-well immune-plates were coated with purified DR5 or *Pfu* DNA polymerase proteins, which was diluted in the coating buffer (NaHCO_3_-NaOH PH 9.6) at the concentration of 1.0 μg/mL and 10.0 μg/mL for 2 h at 4 °C overnight, respectively. 100 μl of supernatants of the nanobody expressing culture media were added to the coated plate for incubation for 2 h at the room temperature and then washed 3 times with the PBST (0.5% v/v Tween 20 in PBS PH 7.4) buffer. As the nanobody were all fused with N terminal His tag. A commercial mouse originated anti-His antibody (1:5000; MerryBio Co) and an anti-mouse IgG-HRP (1:5000;Yeasen) secondary antibody were applied for the detection of the specifical binding nanobody. The reaction was developed with the colorimetric substrate TMB(3,3′,5,5′-Tetramethylbenzidine) and H_2_O_2_ for 15 min, following by the addition of 50 μL 2M H_2_SO_4_ to terminate the reaction. The absorption at 450nm was monitored by a microplate reader (Synergy BioTek H1).

### Isolation of nanobodies

The integrated screening and expression of nanobodies was named the isolation of nanobodies in our work. Firstly, the immunized nanobody library was transformed into the T7 competent cell to obtain a pool of *E. coli* expressing the immunized nanobody library. For the initial screening, a number of 10^n^ transformed *E. coli* were inoculated in a 96 deep-well plate with 2 mL TB medium and 100 μg/mL Amp in each well. A breath-easy membrane was used to seal the 96 deep-well plate. When OD_600_ reached 0.8, 0.5mM IPTG was added for induction overnight at 37 °C. The plate was then centrifuged at 3000 rpm for 30 min to collect the supernatant. To identify the well containing target-specific nanobodies clones, ELISA was performed as aforementioned. A division of λ×10^n^ clones were taken out from the positive well(s), then diluted into 200 mL TB medium and distributed evenly into a second 96 deep-well plate for the next round of target-specific nanobody isolation. The population of positive clone(s) expressing the target-specific nanobody will be enriched by a factor of by a factor of 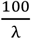 over each round of isolation according to Poisson distribution. The isolation of target-specific nanobody was repeated till λ×10^n^ ≈10. Then, clones from the positive well were spread on the LB agar plate with 100 μg/mL Amp to get separated single clones of *E. coli*. Single clones were then inoculated into a new 96 deep-well plate with one clone in each well for the final round of isolation to get the target-specific monoclonal nanobodies. A value of n in between 1 to 3 was recommended for the initial isolation step dependent on the abundance of the target-specific IgG secreting B cells in the peripheral blood. λ value of 10 was used in this work.

### Isothermal titration calorimetry (ITC)

DR5, *Pfu* DNA polymerase and nanobodies were expressed and purified according to the published protocols ^28,29^. Protein samples were dialyzed to the same buffer (20mM HEPES-NaOH, 150mM NaCl pH 8.0) at 4 °C overnight prior to experiments. Protein concentrations were determined after dialysis. ITC Experiments were performed using the Malvern MicroCal PEAQ-ITC system with 10 to 50 μM antigen proteins or nanobodies in the reaction cell and 100 to 500 μM of the interacting nanobodies and antigen proteins in the injection syringe. Experiments were performed under stirring at 750 r.p.m. at 25 °C. The initial injection was 0.4 μL, following by a series of injections of 2.5 μL. Data was analyzed with the AFFINImeter ^30^ software.

### Nuclear Magnetic Resonance (NMR) Spectroscopy

*Pfu* DNA polymerase and its corresponding monoclonal nanobodies were dialyzed in the sodium phosphate buffer (20mM sodium phosphate,150mM NaCl pH 7.4) at 4 °C overnight. 100 μM of ^15^N-labeled nanobody with 5% of D_2_O were applied for 2D [^15^N,^1^H]-HSQC measurement in the absence and presence of 50 μM *Pfu* DNA polymerase. 50 μM of a mutually different unlabeled nanobody was then added in the NMR sample of ^15^N-labeled nanobody with 50 μM *Pfu* DNA polymerase to identify if two nanobodies shared overlapping binding sites on the *Pfu* DNA polymerase. All NMR spectra were measured at 298 K on a Bruker Avance-600 MHz spectrometer equipped with a TCI cryoprobe. Data were processed with the Topspin software and displayed with the same contour level.

### *Pfu* DNA polymerase activity blocking assay

Real-Time Quantitative PCR was applied to verify the *Pfu* DNA polymerase activity blocking effect of the isolated nanobodies. A single-stranded DNA incubation experiment was performed with 1U *Pfu* DNA polymerase, 200 μM dNTP, 20 mM Tris-HCl, 10 mM KCl, 0.1% Triton-X100, 0.1mg/mL BSA, 2mM MgSO_4_, 10mM (NH_4_)_2_SO_4_, 1X of SYBR Green I, pH 8.8. The experiment group was performed with addition of 0.1μM or 1 μM isolated *Pfu* monoclonal nanobody. The control group was carried on with addition of 0.1μM or 1 μM non-specific nanobody. Another negative control was performed without *Pfu* DNA polymerase. The experimental procedure was 50 °C 1min, 90 cycles. All experiments were run in triplicate.

## Acknowledgments

This work was supported by National Key R&D Program of China 2018YFE0202301, 2018YFE0202300 and National Natural Sciences Foundation of China grants 22174151 and 2199108.

## Author contributions

The research was performed by Zhiqing Tao, Huan Wang, and Xiaoling Zhao; data analysis and validation were performed by Zhiqing Tao; manuscript was written by Lichun He, Zhiqing Tao, Huan Wang, Xiaoling Zhao; design of the experiments and funds acquirement conducted by Lichun He, Xu Zhang, Juan Zhang, and Maili Liu. All authors have given approval to the final version of the manuscript.

