## supplementary data for "A method for rapid nanobody screening with no bias of the library diversity"

### Supporting Figures

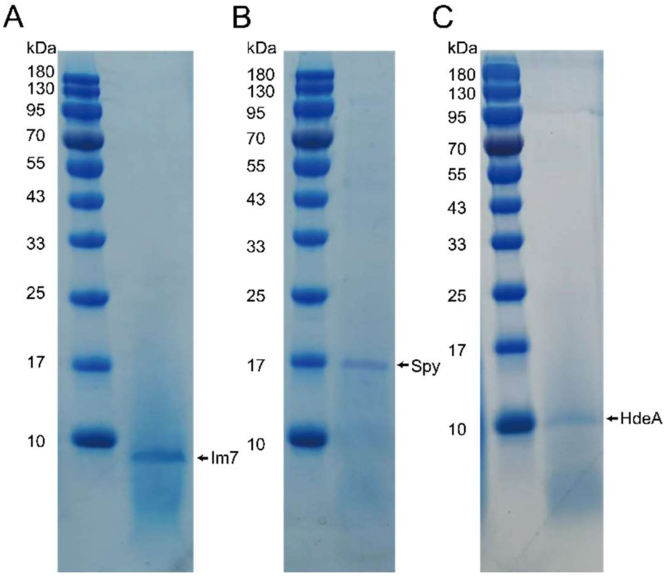


**Supplementary Figure 1.** SDS-PAGE analysis of the secretion of recombinant proteins fused with the pelB signal peptide. A. SDS-PAGE analysis of Im7 protein secreted in the supernatant of the culture medium of *E. coli*. B. SDS-PAGE analysis of Spy protein secreted in the supernatant of the culture medium of *E. coli*. C. SDS-PAGE analysis of HdeA protein secreted in the supernatant of the culture medium of *E. coli*. The supernatant of the cell culture medium was applied for the SDS-PAGE analysis without concentrating.

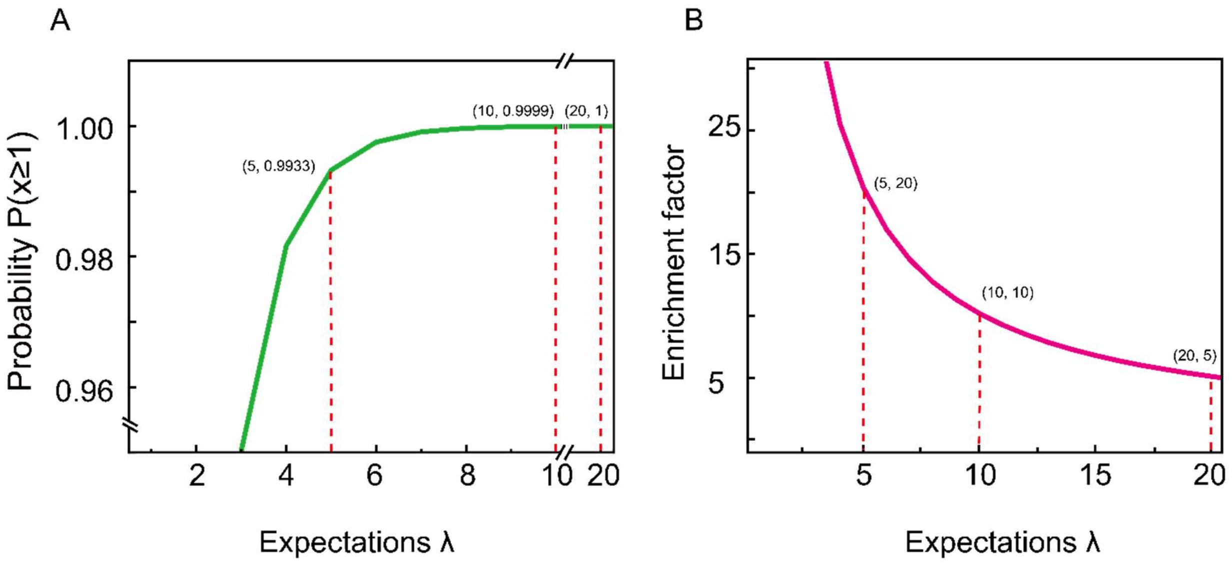


**Supplementary Figure 2.** The theoretical model for the enrichment of positive clones. A. Plot of the probability of at least one positive clone presenting in the division (m) taken from the identified positive well against the value of λ. B. Plot of the enrichment factor against the value of λ.

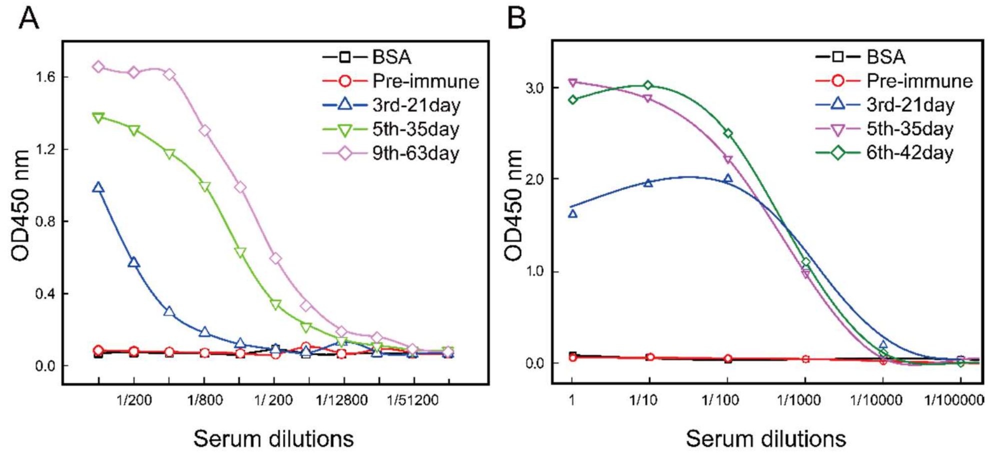


**Supplementary Figure 3.** Assessment of the antibodies in the serum samples from llamas immunized with *Pfu* DNA polymerase (A) and DR5 (B) by ELISA assays. A. secondary rabbit anti-camelid antibody conjugated with the Horseradish peroxidase (HRP) was used for detecting and visualizing the presence of the antigen-specific binding camel antibody.

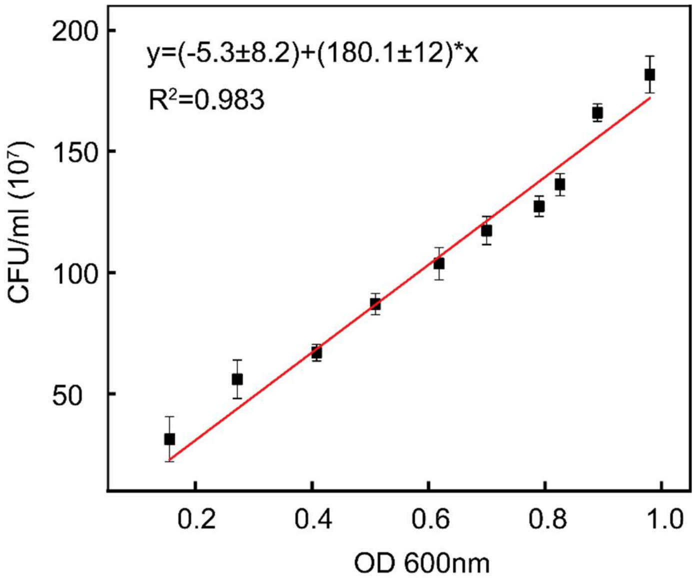


**Supplementary Figure 4.** Standard curve of *E. coli* number against the value of $OD_{600}$. The number of *E. coli* and the value of $OD_{600}$ were graphed on the y-axis and x-axis respectively.

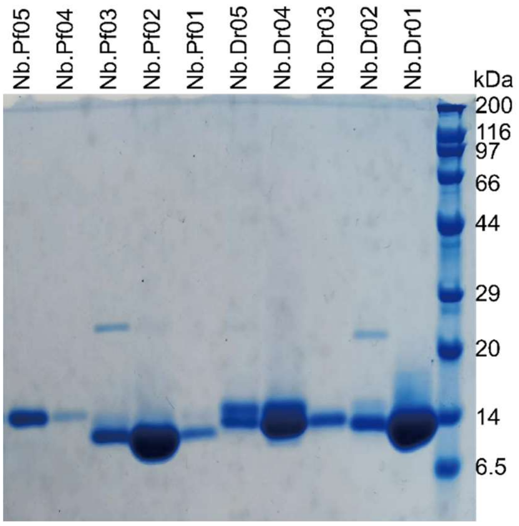


**Supplementary Figure 5.** SDS-PAGE analysis of the elution fraction of each nanobody. 0.1 L supernatant of the cell culture medium of each nanobody was mixed with 1 mL of Ni-NTA beads for 30 mins and then eluted with 5 mL of PBS buffer with 500mM imidazole respectively.

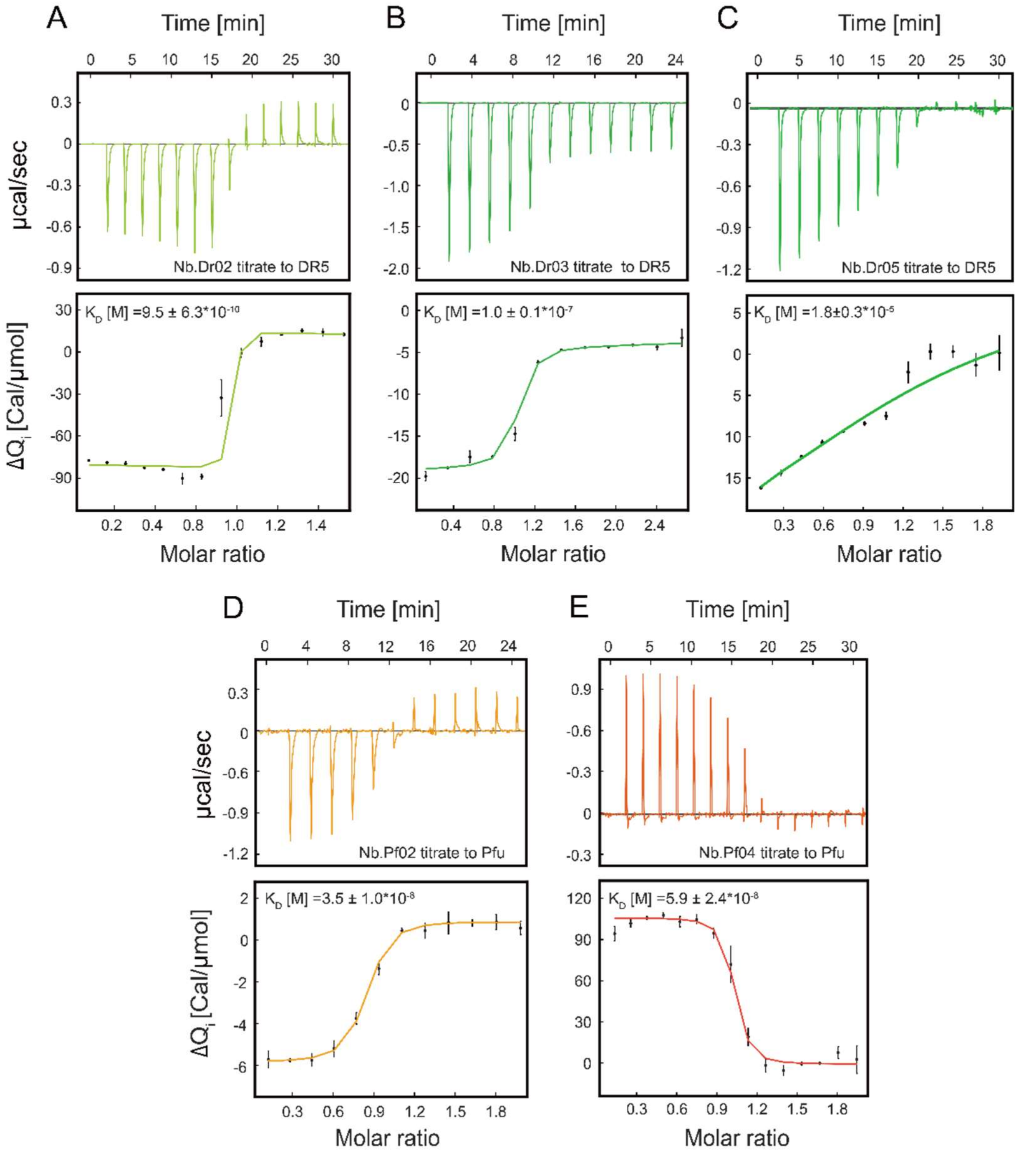


**Supplementary Figure 6.** Determination of the binding affinities of isolated nanobodies. (A-C). Results of ITC measurements of three monoclonal nanobodies Nb.Dr02 (A), Nb.Dr03 (B), and Nb.Dr05 (C) against DR5 protein respectively. (D-E) Results of ITC measurements of two monoclonal nanobodies Nb.Pf02 (D) and Nb.Pf04 (E) against *Pfu* DNA polymerase respectively. Values of the binding equilibrium constant were indicated on corresponding figures.

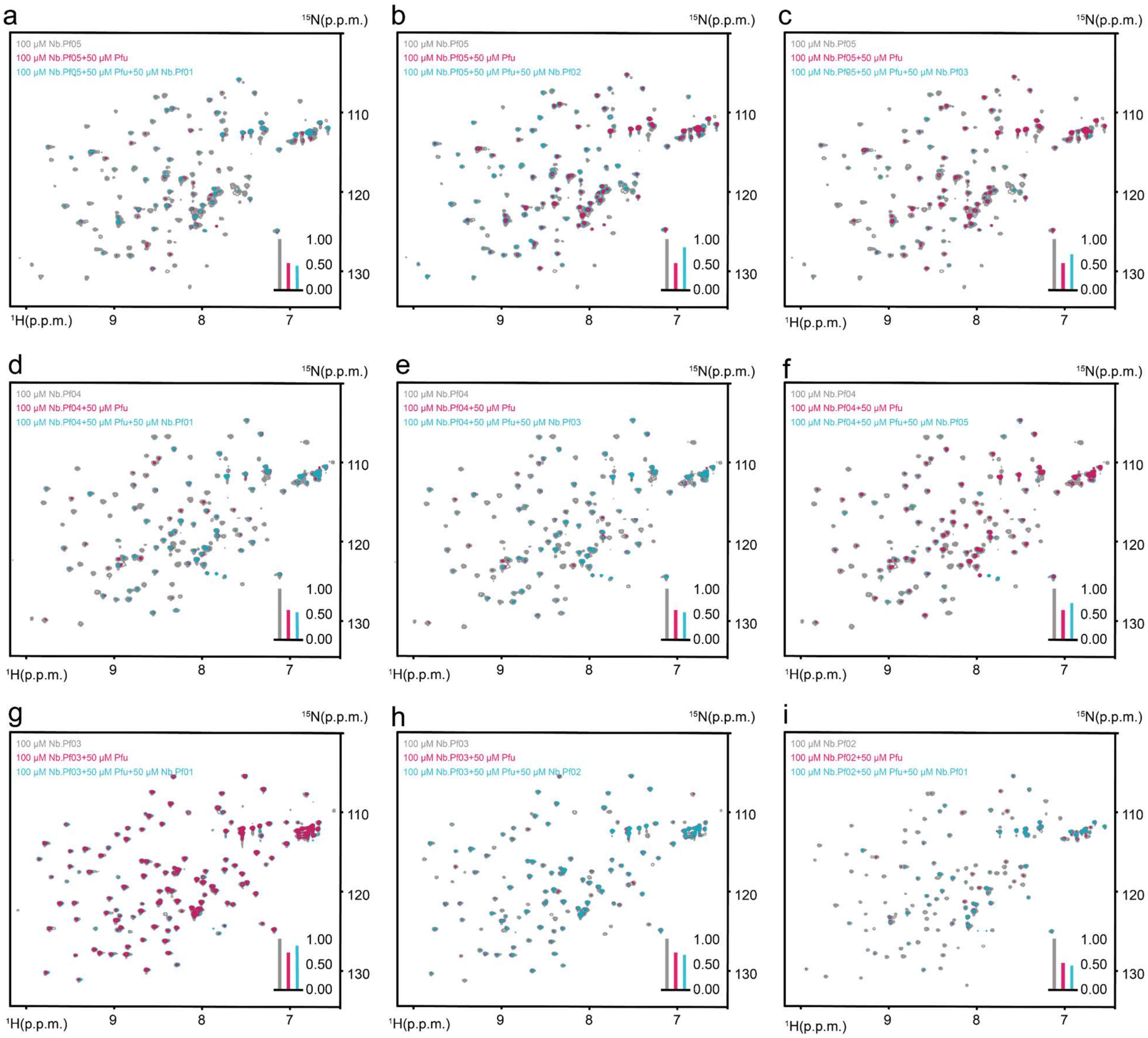


**Supplementary Figure 7.** NMR-based competitive binding assay of different isolated monoclonal nanobodies on *Pfu* DNA polymerase. $^{15}N$-$^{1}H$ HSQC spectra were employed to measure the binding of a $^{15}N$ labeled nanobody with *Pfu* DNA polymerase in the presence of a second unlabeled nanobody. (A-I) The mutual competition statuses of every two isolated *Pfu* DNA polymerase nanobodies were probed. $^{15}N$-$^{1}H$ HSQC spectrum of the $^{15}N$ labeled nanobody was colored in grey. $^{15}N$-$^{1}H$ HSQC spectrum of the $^{15}N$ labeled nanobody in the presence of the *Pfu* DNA polymerase was colored in magenta. $^{15}N$-$^{1}H$ HSQC spectrum of the $^{15}N$ labeled nanobody in the presence of both *Pfu* DNA polymerase and a second unlabeled nanobody was colored in cyan. The concentration of each $^{15}N$ labeled nanobody was 100 μM. The concentration of *Pfu* DNA polymerase was 50 μM. The concentration of the second unlabeled nanobody was 50 μM. The bar graph in the bottom right corner of each spectrum illustrated the relative peak intensity. The gray bar indicated the relative peak intensity of the $^{15}N$-$^{1}H$ HSQC spectrum of $^{15}N$ labeled nanobody. The magenta bar indicated the relative peak intensity of the $^{15}N$-$^{1}H$ HSQC spectrum of $^{15}N$ labeled nanobody in the presence of the *Pfu* DNA polymerase. The cyan bar indicated the relative peak intensity of the $^{15}N$-$^{1}H$ HSQC spectrum of $^{15}N$ labeled nanobody in the presence of both *Pfu* DNA polymerase and a second unlabeled nanobody. All the spectra were recorded at 25 ℃ on Bruker Avance-600 spectrometers. Data were processed in the same way and displayed with the same contour level. and displayed with the same contour level.

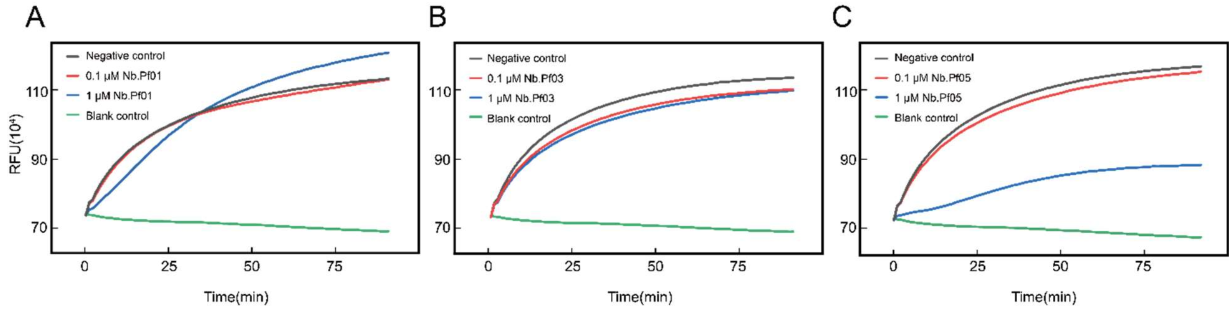


**Supplementary Figure 8.** Assessment of the blocking effect of isolated nanobodies on *Pfu* DNA polymerase. (A-C) Real-Time Quantitative PCR was employed to verify the block effect of Nb.Pf01(A), Nb.Pf03 (B) and Nb.Pf05 (C). The black line represents the control group with *Pfu* DNA polymerase. The red and blue lines represent the experimental group with *Pfu* DNA polymerase and a final concentration of 0.1 μM (red) and 1 μM (blue) of the isolated monoclonal nanobody, respectively. The green line represents the blank group with no *Pfu* DNA polymerase.
